# NeuroVLM: A generative vision-language framework for human neuroimaging

**DOI:** 10.64898/2026.02.06.704508

**Authors:** Ryan Hammonds, Jerjes Aguirre-Chavez, Borngreat Omoma-Edosa, Aarya Patel, Bradley Voytek

**Author notes:** Contributing authors.

## Abstract

Neuroimaging research has produced tens-of-thousands of articles that pair natural language and activation coordinate tables. Recent advances in vision-language models (VLMs) have provided methods to model text and images simultaneously. In this work, we present NeuroVLM, a model architecture for learning from 30,826 human neuroimage-text pairs. The architecture supports contrastive and generative objectives. The contrastive model ranks similarity between neuroimages and text. The generative models include text-to-neuroimage and neuroimage-to-text. These models are evaluated on network images from a variety of atlases, statistical maps from diverse publications, and images created from coordinate tables. These models are capable of generating atlases or maps given a text corpus, generating text interpretations of neuroimages, labeling networks, finding publications most related to a neuroimage query, or finding neuroimages most related to a text query.

## 1 Introduction

Over the past three decades, human functional magnetic resonance imaging (fMRI) neuroimaging research has changed the landscape of our understanding of how the brain gives rise to behavior and cognition. Tens of thousands of experiments have been performed, resulting in a massive, multimodal corpus of peer-reviewed publications that link natural language text to spatial brain coordinates. While powerful, expert decisions on how to interpret these results and which new experiments to run are constrained by one’s own degree of familiarity with the literature. These constraints result in, for example, variations in how individual brain regions are named, or how collections of brain regions are grouped together and named as common networks [1]. Inconsistencies in nomenclature and interpretations in fMRI are exacerbated by small effect sizes [2–4] and large variability in data analysis approaches. Even when working on the same datasets, different research groups can come to different conclusions [5]. Furthermore, many fMRI studies are interpreted in isolation from other human brain maps, such as gene expression, neurotransmitter and modulator distribution, and electrophysiology [6], giving rise to a narrow view of the brain-behavior relationship [7].

Prior to the introduction of modern large language models (LLMs), text-mining of neuroscientific papers was demonstrated to be capable of generating plausible hypotheses by identifying gaps between text concepts mapped to a network graph [8]. In other domains, such as materials science, high-dimensional text embeddings have been shown to be capable of recapitulating concepts such as the periodic table of elements, and complex structure–property relationships in materials [9].

To overcome some of these limitations, a large, diverse training corpus is useful for integrating information across a broad neuroscientific context [7, 10]. This includes diverse datasets that pair text and brain activation maps, enabling models to translate between modalities: text-to-neuroimage and neuroimage-to-text. Past text-to-neuroimage models such as neurosynth [3] and neuroquery [11] use term-frequencies based on a predefined corpus [3, 11]. This limits handling of unseen phrases, synonymy, and context, and assumes near-independence of terms or linear additivity across terms. These methods fail to generalize to out-of-corpus language and to account for non-linear interactions.

Advances in language modeling architectures [12–14] move beyond term-frequencies and instead use transformers to model language. Transformer architectures are standard in LLMs, remove the dependence on a fixed vocabulary, and are context-sensitive. Transformers are also used in vision-language models (VLMs), with the goal of aligning text-image latent vector pairs [15].

Recent works such as neurocontext [16] and niclip [17] use LLMs to encode text from neuroimaging publications. Contrastive loss [15] was used in these studies to learn a bi-modal embedding space, aligning text and neuroimage representations. These models provide similarity-based ranking and retrieval, given a corpus of publications and neuroimages. While powerful, contrastive models cannot generate brain images from input text, nor text outputs from 3D brain maps.

Here we present, NeuroVLM (Fig. 1a), a lightweight, artificial intelligence (AI) framework that trains both generative and contrastive models. This framework embeds text-neuroimage pairs to a shared latent space. This latent space can be trained with a contrastive objective to support ranking and retrieval, which allows us to label neuroimages, e.g., spatial ICA components, with text, and to select neuroimages most related to a text query. Decoding the latent space provides generative text-to-neuroimage and neuroimage-to-text models.

**Fig. 1.**
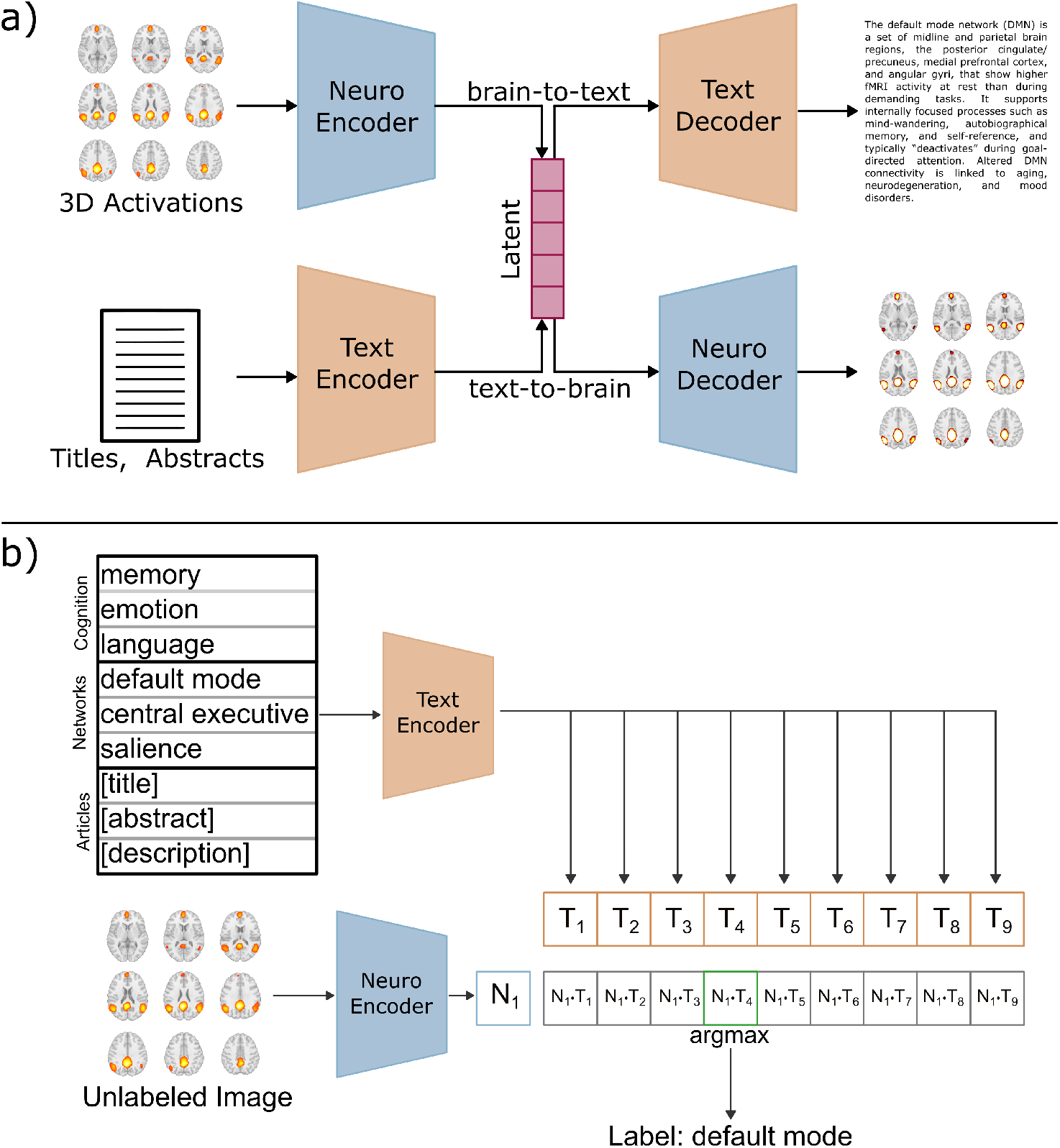
Model architecture. **a)** NeuroVLM maps pairs of activation maps and text, from the same publication, to a shared latent vector (pink). Decoding the latent vector results in brain-to-text or text-to-brain generation. **b)** The shared latent vector is the basis for contrastive models, e.g., ranking and retrieving text most similar to a query brain map. Given an unlabeled, query activation map, the method ranks how similar a corpus of text is to the image. All encoded vectors, *N*_1_ and {*T*_1_, …, *T*_9_} are unit normalized. The inner product between these vector pairs is cosine similarity. Taking argmax gives the most similar text. Any text corpus or image set may be used, suitable for a variety of use cases. The direction may also be flipped from what is presented: comparing a single text query to a set of activation maps.

To improve the accessibility of these models, we significantly reduced the computational requirements by using much smaller text encoders [18] with 110 million parameters, rather than depending on LLMs with tens-to hundreds-of-billions of parameters. Each component of NeuroVLM is modular and interchangeable with larger or smaller models depending on requirements.

The resulting open-source models support a variety of applications. Use cases include generating atlases or maps from a text corpus or query, which could be used for first-pass hypothesis generation. NeuroVLM can also generate natural-language interpretations of human brain maps (Fig. 2) or automatically label functional networks (Fig. 5b). Finally, it supports cross-modal retrieval by finding the publications most related to a statistical map, or the statistical maps most related to a text query.

**Fig. 2.**
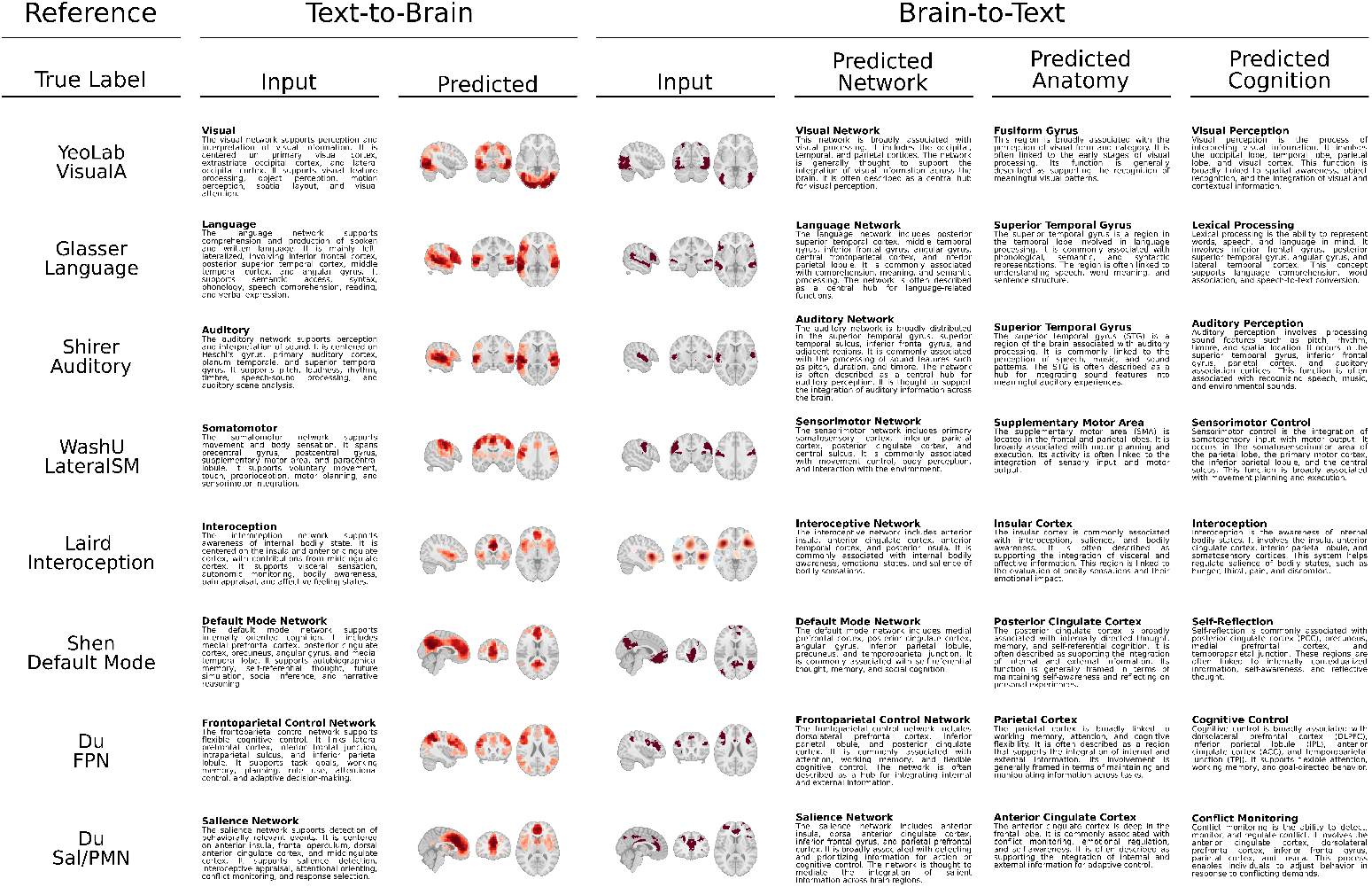
Networks: text-to-brain and brain-to-text generation. The input to text-to-brain generation was a high-level summary of the reference image. The input to brain-to-text generation were atlas networks, see the reference column for the group who produced the map. Text-to-brain generation used text description of networks to generate a set of activation maps. Brain-to-text generation used atlas images to generate interpretations of the brain map. Network, anatomy, and functional tokens were stacked into token embedding matrix, predicted from the Qformer. This provides steering of brain-to-text interpretation to corresponding aspects.

## 2 Methods

### 2.1 Datasets

Data were aggregated from PubMed central, neurosynth compose [19], the cognitive atlas [20], the network correspondence toolbox [1], and NeuroVault [21]. The networks toolbox maps [1], NeuroVault maps [21], and a subset of the PubMed articles were left out as test sets. The cognitive atlas is a dataset of text labels used to label activation maps post-training.

1. PubMed Central: 30,826 publications with titles, abstracts, and coordinate tables
2. Cognitive Atlas [20]: 1,000 cognitive terms and definitions
3. Networks [1]: 100 labeled network images
4. NeuroVault [21]: 3,000 group-level activation maps, titles, and abstracts

### 2.2 Preprocessing

Coordinates were convolved in MNI space with a 9 mm activation Gaussian kernel, following [11]. The resulting activation maps were transformed using the dictionaries of functional modes (DiFuMo) atlas [22]. Reconstruction replaces voxel values with a weighted combination of DiFuMo components. Component coefficients were computed as the DiFuMo-weighted mean activation and were used to reconstruct voxel space by re-expanding the coefficients through the DiFuMo voxel maps. This returns a voxel-space map from a linear combination of DiFuMo components (see Sec. 6.1 for details).

### 2.3 Training

Training was performed with 30,826 text-neuroimage pairs, primarily from PubMed. Text included titles and abstracts of publications. Neuroimages are defined as sparsely clustered, MNI-space activation maps. PubMed training articles instead provide coordinate tables, which were smoothed in 3D, MNI space, similar to the approach used in Neuroquery [11]. Continuous statistical maps are provided in the network and neurovault datasets; these were used as additional test sets. DOIs used in the PubMed training set were excluded from all test sets.

Software, including training scripts, are publicly available at:

https://github.com/neurovlm/neurovlm

Pre-trained models and datasets are available on HuggingFace:

https://huggingface.co/neurovlm

### 2.4 Model Architecture

Building upon past work, we outline a general vision-language framework for neuroimaging applications: NeuroVLM (Fig. 1a). The encoders in this framework map from high-dimensional text and activation-map space to low-dimensional latent vectors (Fig. 1a, pink). After training, pairs of text and neuroimages encode to highly similar vectors. Decoders are able to generate text or activation maps from latent vector embeddings. These encoders and decoders collectively provide text-to-image and image-to-text generation. This framework is general and does not make assumptions on the choice of encoder or decoder architectures.

### 2.5 Autoencoder

An autoencoder was trained to compress 3D activation maps to 384-dimensional latent vectors (Fig. 1a, blue). Maps were down-sampled, masked, flattened, and binarized before passing through fully-connected linear layers with rectified linear unit (ReLU) activations. Exact model architecture, including layer sizes and activation functions, are provided in Table S1. The downsampling and masking procedure was adapted from [11] and moves from a 1 million dimensional MNI space to 28,542 dimensional space. The encoder transforms from 28,542 to a 384 dimensional latent representation. A key component to successful training was the binarization of the activation map prior to training and the use of binary cross entropy loss. Any continuous map, e.g., a z-score or p-value map, can be binarized with an appropriate threshold and clustering method. Another key component in training was sparsity, reducing the influence of dense maps that were fully activated and lacked sparse clusters of activation.

### 2.6 Text Encoder

NeuroVLM uses Specter [18], a small, efficient, and accessible transformer. This small model allows deploying the model on edge devices with little computational requirements. Specter is based on SciBERT [23] and incorporates citation-based connectivity graphs. Specter encodes title-abstract pairs to 768-dimensional latent vectors. Despite being trained on title-abstract pairs, Specter has an ad hoc query adapter, suitable for short form user queries.

### 2.7 Text Decoder

Brain-to-text generation requires a language model. We trained Qwen3-0.6B on 1.2 million neuroscience articles and used this to generate descriptions of images using a query transformer (Qformer) [24]. The Qformer learns a mapping from the latent image space to the token embedding space of LLM. This training was supervised such that the decoded text was optimized to the true, paired text.

The Qformer receives two complementary inputs: a raw 384-dimensional image latent from the image encoder and a 384-dimensional semantic latent, from the contrastive model. Each input is projected into a shared 512-dimensional hidden space, producing two memory tokens: one preserving the raw-image latent signal and one for semantic alignment information. A set of 32 learned query embeddings then attends to these memory tokens through a 6-layer transformer decoder. The resulting query states are mapped into the 1024-dimensional embedding space of the language model, yielding 32 continuous prompt tokens for downstream text generation.

At inference, the images may come from a variety of distributions, including binary masks and t-, z-, F-, and p-maps. The one requirement of the brain-to-text model is to transform these maps to the range: [0, 1]. In addition to a variety of map distributions, thresholding and clustering depends on a variety of parameter choices. To help improve generalization of brain-to-text generation for out-of-distribution maps, relative to the training maps, we introduce a canonical, or stereotyped, projection matrix. This canonical projection matrix is constructed from clean examples of networks, regions, and cognitive concepts. Vector projection produces a denoised semantic conditioning vector constrained to lie near interpretable prototypes. The canonical projection adds a layer of interpretability and can warn if the input image lies too far from known images. See supplementary section S9 for details.

### 2.8 Contrastive Learning

Contrastive learning maximizes cosine similarity between true text-image pairs and minimizes similarity between mismatched pairs [15, 17]. After training, unlabeled images can be labeled by ranking cosine similarity between the target image and any text corpus (Fig. 1b). Pretrained encoders can be used to label neuroimages with networks [1], cognitive concepts [20], or publication descriptions. Inversely, a set of images may be ranked in similarity, relative to target text.

### 2.9 Generative Models

#### 2.9.1 Brain-to-Text: Generating Interpretations

A Qformer [24] is used to generate descriptions of neuroimages (Fig. S1a). The Qformer transforms from latent image representations to the token embedding matrix. The token embedding matrix is decoded by a fine-tuned neuroscience-focused version of Qwen3-0.6B [25].

Training data included image-(title, abstracts). Text was summarized to general text, removing study-specific details. This allows the model to generate higher-level descriptions of activation maps. A final fine-tuning dataset was generated using the text-to-brain model and synthetic descriptions of common networks, regions, and cognition. This step allowed more control of the style of the LLM output, pushing outputs towards shorter, general descriptions. The target was concise brain-to-text captions, rather than plausible abstracts. Examples of brain-to-text generation are provided (Figures 2 and 3).

**Fig. 3.**
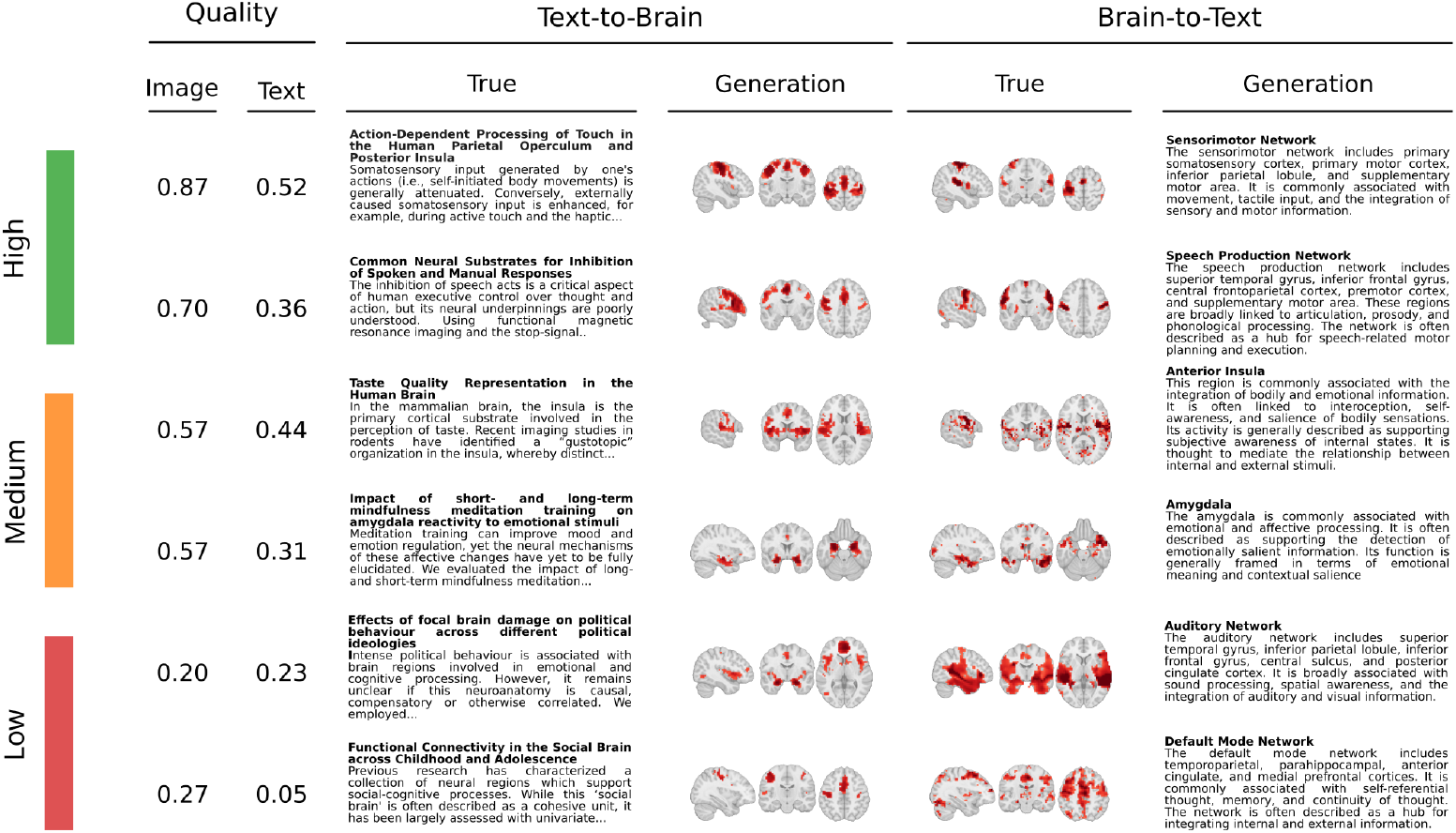
NeuroVault: text-to-brain and brain-to-text generation. Pairs of group-level activation maps and publication text (titles and abstracts) were taken from NeuroVault. Text was used to generate brain maps (left) and true brain maps were used to generate related text (right). Quality of generations is measured with cosine similarity between true and generated.

#### 2.9.2 Text-to-Brain: Generating Activation Maps

Activation maps are generated by projecting text embeddings to the latent space of the neuroimage autoencoder (Fig S1) and using the neuroimage decoder (Fig. S1b). Specter2 [26] is used to embed arbitrary text: titles, abstracts, descriptions, or user queries. A small neural network, the projection head, maps these embeddings to the shared latent space. The projection head allows the language model weights to be frozen while providing fine-tuning, similar to low-rank adaptation (LoRA) adapters [27]. The neuroimage decoder then transforms the latent embeddings to MNI space.

### 2.10 Performance Metrics

The main performance metric used for test evaluation was the recall@k curve. This measure was used for both contrastive and generative models. Cosine similarity between all pairs of latent text and neuroimages is computed. Recall@k measures how often the true neuroimage-text pair was found in the top-k most similar pairs [16]. Given a reference latent neuroimage, recall@k measures how often the paired latent text is in the top-k most similar pairs. The inverse is performed using a reference latent text and computing recall@k against latent neuroimages. We step k from 1 to the max size of the test set and compute recall@k for each threshold. The area under this curve is reported, where 0.5 is chance performance and 1.0 is perfect performance. Recall@k is used for the contrastive model, examining how well aligned the text and neuroimage representations are. For generative models, recall@k is computed from latent representations of generated outputs and true targets.

In addition to recall@k, we also provided contrastive cosine similarity, Dice score, Pearson correlation, and spin-tests for text-to-brain generation. For brain-to-text generation, contrastive cosine similarity, BERTScore, semantic cosine similarity, and a contrastive null test were performed. Matched distributions, or true (image, text) pairs, were compared against a randomly shuffled distribution of (image, text) pairs (Fig. 4).

**Fig. 4.**
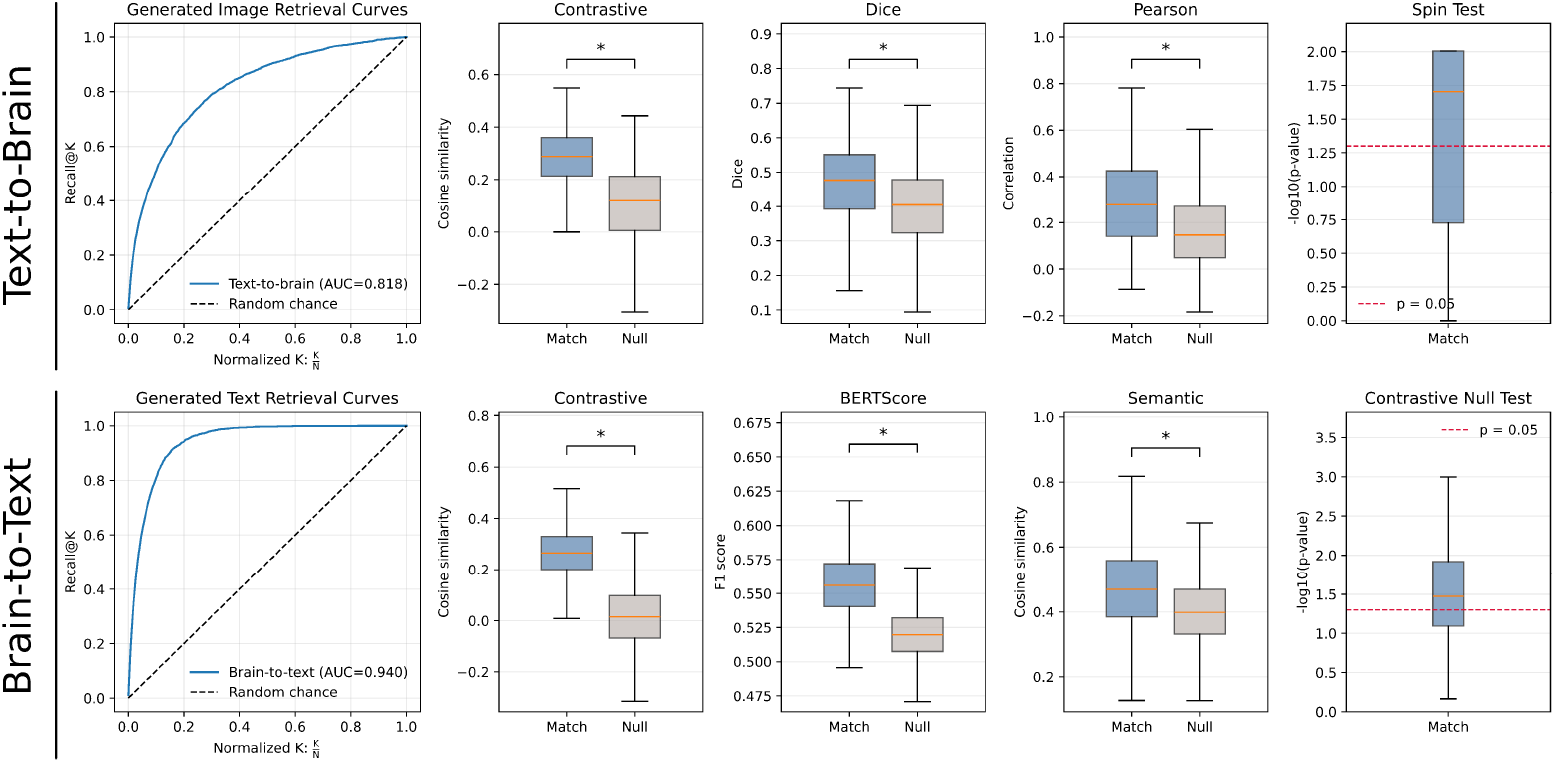
Generative Performance. Text-to-brain and brain-to-text generation performance on the PubMed test set (*n* = 2987). Top row: text-to-brain generation was evaluated by normalized recall@k retrieval, contrastive cosine similarity, Dice overlap, Pearson correlation, and spin-test significance. Bottom row: brain-to-text generation was evaluated by normalized recall@k retrieval, contrastive cosine similarity, BERTScore F1, semantic cosine similarity, and a contrastive null test. Matched pairs are compared against mismatched null pairs where applicable; recall AUC ranges from random chance (0.5) to perfect retrieval (1.0). Null distributions are in gray, they represent randomly shuffled pairs. T-tests were conducted to compare all match versus null distributions, ^∗^*p <* 1*e −* 100 was found for all comparisons.

## 3 Results

### 3.1 Generative Models

#### 3.1.1 Networks

The network dataset [1] was used to demonstrate text-to-brain and brain-to-text generative models, because it provides some degree of ground truth, given that the collection of networks were labeled by experts (Fig. 2 for examples, Fig. 5b for quantitative metrics). The text-to-brain model transformed networks’ definitions to the latent space using the encoder. The latent vectors were then transformed to neuroimages using the decoder. The brain-to-text model generates text related to a neuroimage input from the network dataset. This brain-to-text model uses a Qformer to map from latent image space to the latent token embedding space [24] (Fig. S1).

#### 3.1.2 NeuroVault

Generative neuroimage-to-text and text-to-neuroimage models were evaluated on the neurovault test set, which provides unthresholded brain maps not in the PubMed training set. Qualitative results are shown in Fig. 3, where each row in this figure corresponds to one group-level activation map and study from neurovault. Titles and abstracts from neurovault were used to generate 3D brain maps; only true and predicted (generated) titles are shown.

#### 3.1.3 PubMed

The PubMed test set (*n* = 2987) was used to evaluate both generative directions (Fig. 4). Retrieval performance was quantified with a normalized recall@k curve. For each generated output, we computed its similarity to every candidate item in the test set using the NeuroVLM contrastive embedding space, then asked whether the correct paired item appeared within the top-*k* ranked candidates. For text-to-brain generation, generated brain maps were embedded with the image contrastive projection head and ranked against the source text embeddings. For brain-to-text generation, generated text was embedded with the text projection head and ranked against the source image embeddings. We report the area under the recall curve over normalized rank, 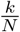, so that random retrieval has expected AUC 0.5, while perfect retrieval has AUC 1.0.

Perfect performance would mean that every generated output is most similar to its true paired input than to any mismatched item in the test set.

In addition to retrieval AUC, we evaluated the paired contrastive cosine similarity between each generated output and its true conditioning input. This metric directly measures alignment in the learned NeuroVLM multimodal space. Matched pairs were compared against a null distribution formed by mismatching generated outputs with inputs from other PubMed examples. Thus, higher matched cosine similarity indicates that the generated brain map or generated text preserves information that is specifically aligned with the correct paired modality, rather than only matching broad dataset-level structure.

For text-to-brain generation, we also quantified spatial accuracy of the generated brain maps. Dice score measured overlap between thresholded predicted and true activation maps. Pearson correlation measured voxelwise spatial correspondence between predicted and true maps, capturing graded similarity beyond binary overlap. Finally, spin-test p-values assessed whether the observed spatial overlap exceeded what would be expected under spatially autocorrelated cortical null maps. The spin-tests were limited to examples that had predominantly surface activation. This provides a more conservative test of anatomical specificity, since neighboring brain locations are not statistically independent.

For brain-to-text generation, we evaluated the generated descriptions using complementary text-based and contrastive metrics. BERTScore F1 measured contextual token-level overlap between generated and reference PubMed text, while semantic cosine similarity measured sentence-level semantic agreement using text embeddings from a generic language embedding model, `all-MiniLM-L6-v2`. Because reference PubMed abstracts can contain broad methodological and domain-general content, these lexical and semantic metrics may remain high even for mismatched pairs. We also included a contrastive null test: for each generated text, we compared its NeuroVLM contrastive similarity to the true source brain map against its similarity to mismatched brain maps from the test set. The resulting empirical p-value measures whether the generated text is more specifically aligned to its own source brain map than to other plausible PubMed brain maps.

### 3.2 Contrastive Model

#### 3.2.1 Network Labeling

Cosine similarity between text-image embeddings was used to label 300 network images [1]. The argmax of cosine similarity is taken as the predicted label. The confusion matrix shows a vast majority of images were labeled correctly using this approach (Fig. 5b). Network images were embedded to a common latent space and then compared against latent text embeddings corresponding to networks, regions, and cognition. Example network images and associated text are shown in Fig. 2.

The network dataset [1] also contains unlabeled independent components (IC) for the Human Connectome Project and UK Biobank. Independent component analysis (ICA) of resting state networks has historically required manual classification of networks versus noise components [28]. Semi-supervised approaches also require manual classification for a classifier to learn from [29, 30]. These methods classify components as either noise or target components. The ICA components from the network dataset are labeled automatically using the contrastive model Fig. S4, with more detail, e.g., the names of various networks. Although noise is not directly modeled, noise components will have low similarity with a resting-state text corpus.

#### 3.2.2 NeuroVault

The NeuroVault [21] test set was used to test the contrastive model. This shows that the model generalizes from coordinate tables during training to group-level activation maps at inference. Recall@k measures how often the true neuroimage-text pair was found in the top-k most similar pairs (Fig. 5b, left) [16]. Similar to AUROC, the area under the curve is expected to be 0.5 for a randomly initialized, untrained model. After training with the PubMed set, the AUC reached 0.85.

#### 3.2.3 PubMed

The PubMed test set was used to test the contrastive and generative models (Fig. 5a). These results are from 5-fold cross-validation, showing moderate AUC for the contrastive models (AUC=0.82). Additional results show that the contrastive model performs similarly to neurocontext and niclip (Fig. S3), with AUC ranging from 0.80 to 0.83. All test examples in this comparison were left out of training from all three models.

### 3.3 Autoencoder

The first step in training is to learn an autoencoder that can reconstruct neuroimages. This was performed on the PubMed training set. Each MNI space activation map is encoded to a 384-dimensional latent vector then decoded back to MNI space. This latent vector space is the target for encoding text, resulting in text-to-brain decoding. The autoencoder was trained on binarized targets and predicts logits, e.g., each voxel was modeled as a binary classification problem. The model output is interpreted as the probability of each voxel being active. Performance is quantified by area under the receiver operating characteristic (AUROC = 0.975) and normalized negative log likelihood (bits/voxel) [31] that measures improvement relative to a naive baseline (Fig. 5d). The test set included network maps, NeuroVault images, and PubMed (Fig. 6).

**Fig. 5.**
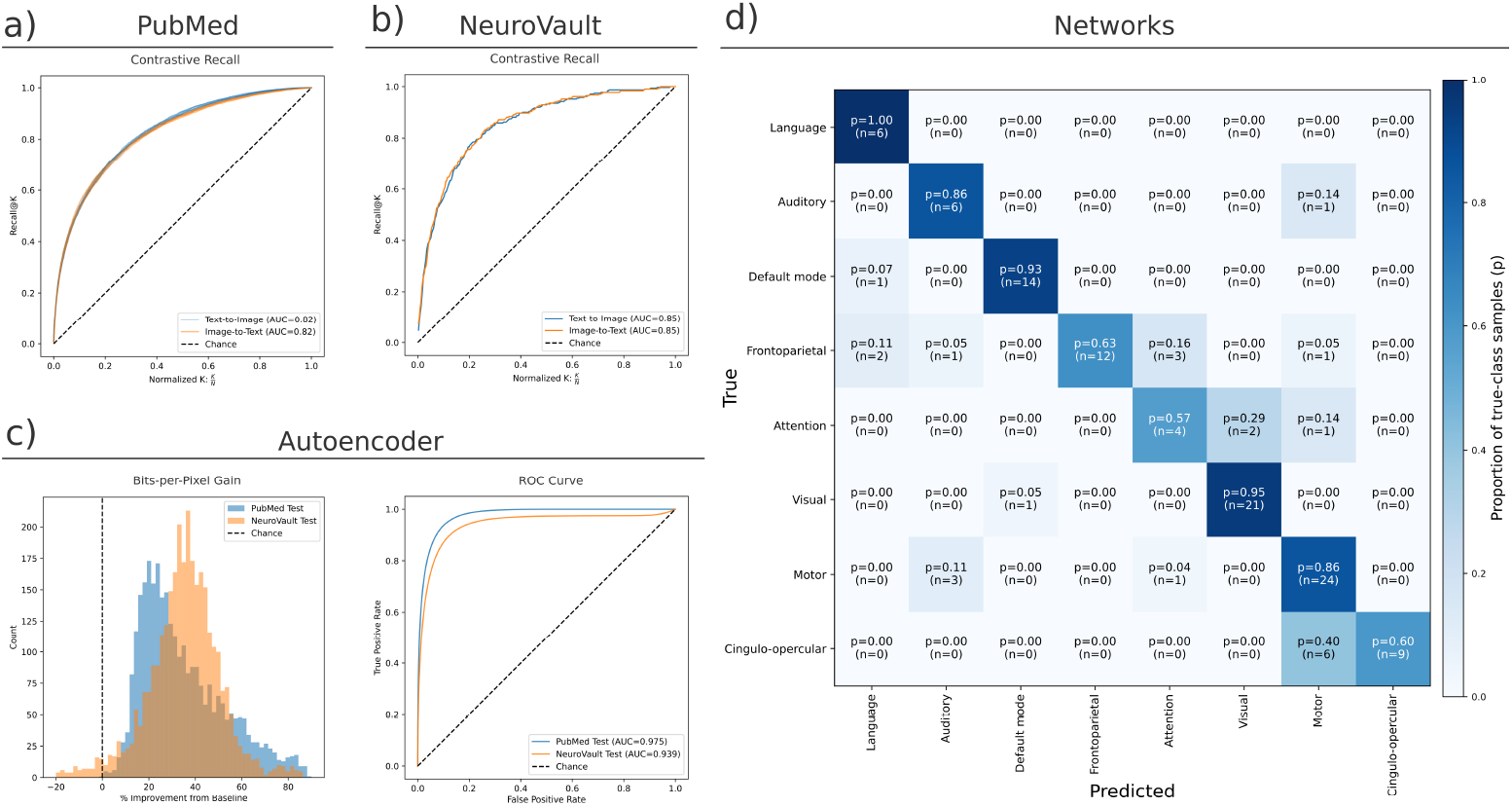
Contrastive Performance. Results are presented for the neuroimage autoencoder, contrastive models, and generative models. Recall@*k* has expected random performance of area-under-the-curve (AUC=0.5), along the diagonal. Perfect performance is AUC=1.0. Recall@*k* measures how often the true pair is in the top-*k* most cosine similar pairs. **a) PubMed**. Contrastive recall curves (AUC=0.81). **b) NeuroVault**. Contrastive recall (AUC=0.78) curve (AUC=0.85). **c) Autoen-coder**. Binary voxel performance (AUCROC=0.94) and bits-per-pixel gain from a naive baseline. **d) Network classification**. Multi-class predictions, from argmax of cosine similarity between contrastive embeddings, as a confusion matrix. The diagonal is correct classifications (dark blue). Counts and proportion of true-sample class are given per cell.

**Fig. 6.**
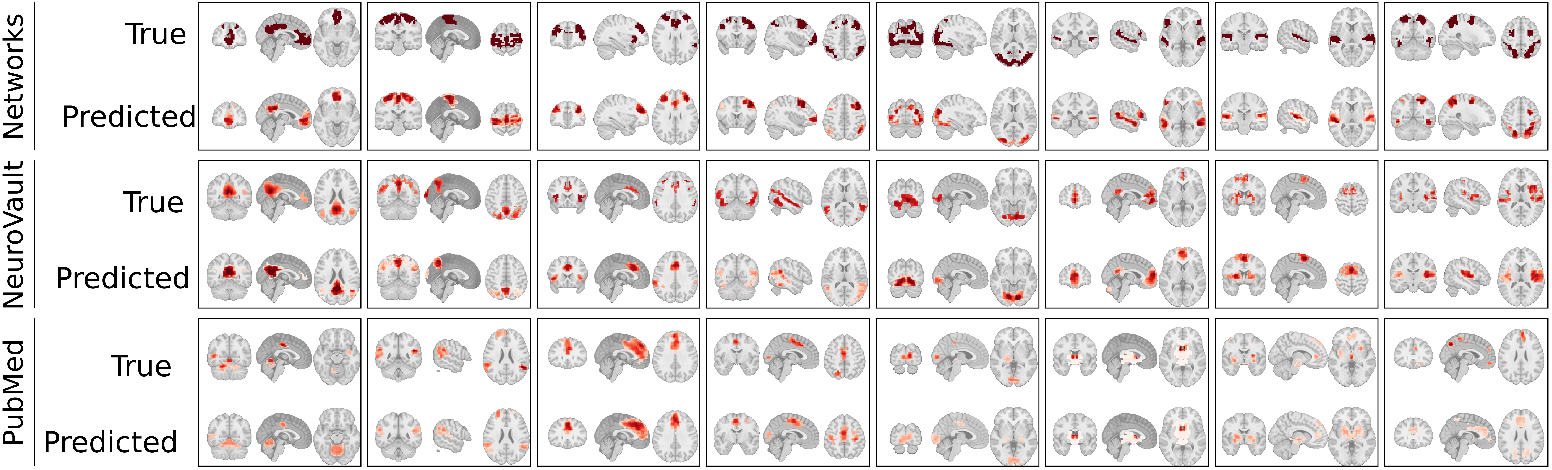
Autoencoder test set reconstructions. Example reconstructions for the networks, NeuroVault, and PubMed datasets. The top in each box is the true image. The autoencoder encodes the images to latent space, then decodes the latent space to recreate the image. The reconstructed predictions are shown below each true image. All images were randomly selected from the test sets.

## 4 Discussion

One of the most central dogmas of neuroscience is that of functional localization: the idea that specific cognitive and behavioral functions can be localized to specific brain regions. This view has been updated to a richer, network-level view of behavior and cognition. The history of human brain imaging is such that vast amounts of natural language data have been generated in attempts to describe the populations from whom brain images were recorded, under what experimental conditions, with expert narratives describing the resulting brain maps. Together, these texts and images provide a rich set of data upon which we collectively build novel hypotheses about the neural basis of behavior, cognition, and disease.

Here, we introduce NeuroVLM, a generative AI framework that can serve as an expert collaborator in this process of narrative building and hypothesis generation. This generative approach builds upon past works that focused on text-to-brain models, constrained to a finite, predefined corpus [3, 11]. The text-to-brain model was expanded to BERT-based text encoders [32]. More recent efforts have trained contrastive models to rank similarity between text-brain pairs using LLM embeddings [16, 17].

NeuroVLM is an integrative framework (Fig. 1a) that is capable of generating text from neuroimages or neuroimages from text. This framework relies on encoding to, and decoding from, a common latent space. This general framework is not dependent on the choice of encoders or decoders. Instead, we provide a neuroimage autoencoder, a model to project text embeddings to the shared latent space, and an interpretable method to generate text from neuroimages. These models were trained on 30,826 publications and extend to group-level statistical maps [21] and network atlases [1]. Use cases include generating atlases or maps given a text corpus, generating text interpretations of 3D brain maps, labeling networks with coactive brain regions, finding publications most related to a statistical map query, or finding statistical maps most related to a text query.

A central limitation to our approach is that our training neuroimages are not native statistical maps, but proxy maps reconstructed from reported peak coordinates and smoothed with a fixed kernel. Coordinate tables are not intended to be a sufficient statistic for the underlying effects –– they omit subthreshold structure, cluster shape, effect-size information, variance, and many study-specific details such as contrast definitions, preprocessing choices, and sample characteristics. This choice is pragmatic for scale, but it bounds the quality of what the model can learn.

Another limitation involves topics that are under-powered and under-studied. While common networks and widely used terminology are well-represented, rare constructs, emerging topics, or underrepresented populations can be mapped less reliably. For more niche, specific topics, additional fine-tuning is recommended.

In addition, it is important to emphasize that generative models often can introduce confident predictions that go beyond the evidence. Using NeuroVLM, brain maps will be generated from any text input, no matter how ludicrous. For example, 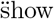 me the hot dog areas of the brain, 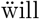 result in beautiful 3D brain images – largely regions related to gustation and reward, as it turns out (so perhaps not entirely ludicrous). Nevertheless, NeuroVLM should be viewed as an AI collaborator – a tool for organizing and navigating the literature, suggesting candidate networks, concepts, and related papers – rather than as an oracle that provides mechanistic explanations or ground truth maps.

Despite these caveats, our work suggests that large-scale, paired text-neuroimage corpora are sufficient to learn a useful shared representation that supports practical tasks such as retrieval, labeling, and coarse generation. Our strongest limitations involve access to neuroimaging data rather than model architecture. Contributions to open-access repositories, such as NeuroVault [21] and OpenNeuro [33], will improve the future generations of multimodal neuroscience models; or more simply: higher quality, expertly labeled and annotated data can yield better results [34].

Looking ahead, we believe that it is critical to strengthen the ecosystem around standardized benchmarks and reproducible evaluation. Network labeling and recall-style retrieval provide accessible demonstrations, but they do not fully measure neuroscientific correctness, robustness to analysis pipelines, or calibration under dataset shift. Progress will benefit from community benchmarks that specify train-test splits, map types, preprocessing, and query sets, and that evaluate multiple axes of performance: retrieval utility, interpretability, out-of-distribution generalization, and failure modes. Open model checkpoints, transparent preprocessing, and reproducible training scripts should be prioritized.

NeuroVLM frames the neuroimaging literature as a bidirectional link between how studies are described in language and where effects are reported in the brain. This enables a single framework to support retrieval, labeling, and generation from tens-of-thousands of paired examples. These models can help organize a fragmented literature into a searchable, cross-modal interface: a way to move from maps to candidate concepts and related papers, and from text queries to spatial priors and exemplar results. By making these capabilities accessible with modular components and modest compute, this work reduces the barrier to using modern multimodal methods for everyday neuroscientific workflows.

Our results underscore that the limiting factor for biological fidelity is often the structure of what the field reports and shares. When the community primarily provides peak coordinates, models inherit the constraints of peak-coordinate reconstructions; in contrast, sharing more data, such as full statistical maps, contrast definitions, and rich metadata, creates a virtuous cycle and allows models to learn quantitative, uncertainty-aware representations that better reflect the evidence [34].

NeuroVLM is a step toward computational tools that augment, rather than replace, expert interpretation. Its most immediate value is as a literature navigation and hypothesis-generation aid that can surface relevant work, highlight commonalities across maps and descriptions, and provide interpretable suggestions. The broader opportunity is to couple these models with standardized benchmarks and richer open data so that future systems are reliable, reproducible, and grounded in the full statistical record of human neuroimaging research.

## Supplementary information

## 5 Generation

**Fig. S1.**
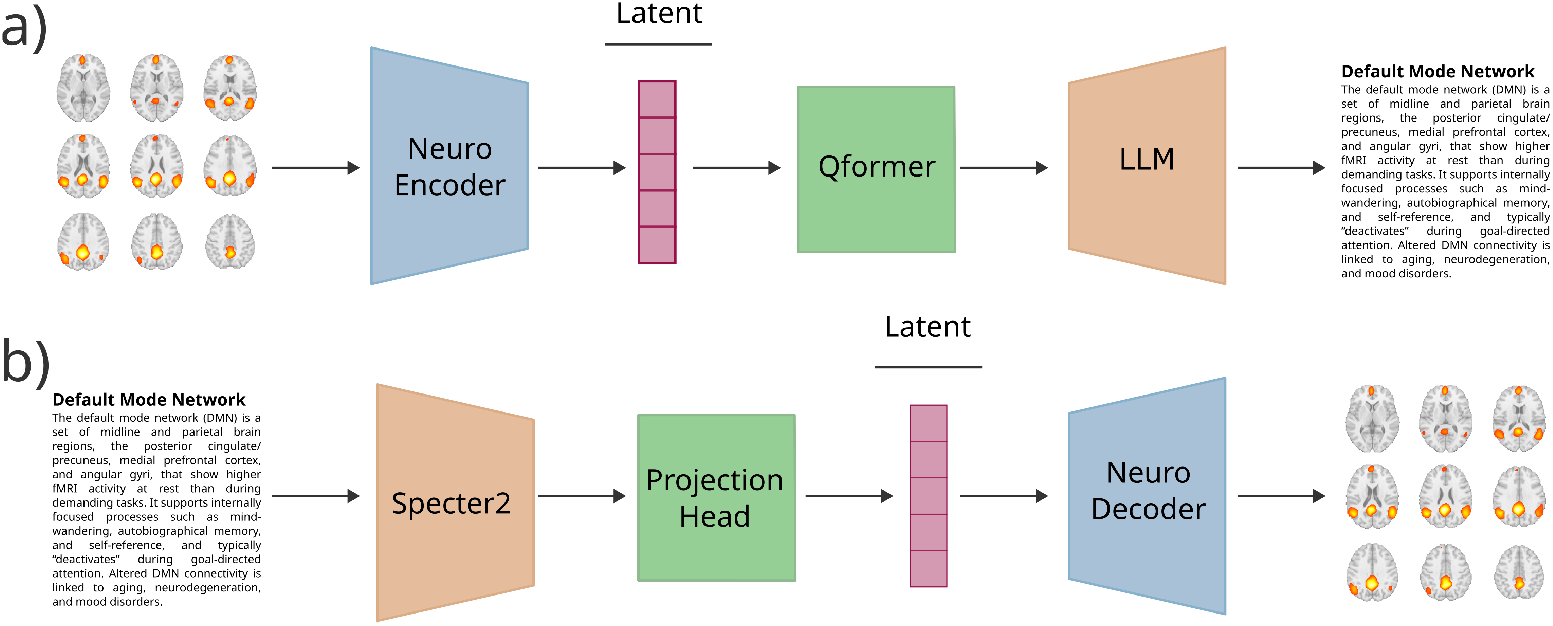
Generating and interpreting neuroimages. **a)** Brain-to-text generation is used to predict text interpretations of input brain maps. An activation map is first encoded to the latent vector (pink). The Qformer (green) then transforms the latent vector to the token embedding matrix of the LLM. Qwen3-0.6B (orange) generates text descriptions. **b)** A text query is first encoded with Specter2, a frozen, pre-trained text encoder. The projection head is a small, trainable network that transforms from Specter2’s embedding space to the bi-modal, text-neuroimage embedding space (pink). This latent vector is decoded (blue) to generate activation maps.

## 6 Preprocessing

### 6.1 Coordinate Tables

Coordinate tables (n=30826) were collected from PubMed and NeuroSynth Compose databases. Coordinates were convolved with a 9mm Gaussian kernel. The kernel was normalized to values between zero and one. The zero-to-one range allows using binary cross-entropy loss when training the autoencoder. The MNI-space convolutional product, **X**, was clamped to values between zero and one. The 512-dimensional DiFuMo atlas, **P**, was used to transform the convolved products (Fig. S2). This is in contrast to using the 512-dimensional embedding directly, as in [16, 17]. Instead, the autoencoder is trained to learn a data-driven, low-dimensional embedding space.

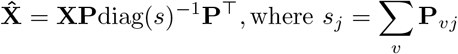

The DiFuMo reconstructions were resampled to 4mm isotropic voxel space, matching [11]. Masking and flattening produced a 28,542 dimensional vector per image.

**Fig. S2.**
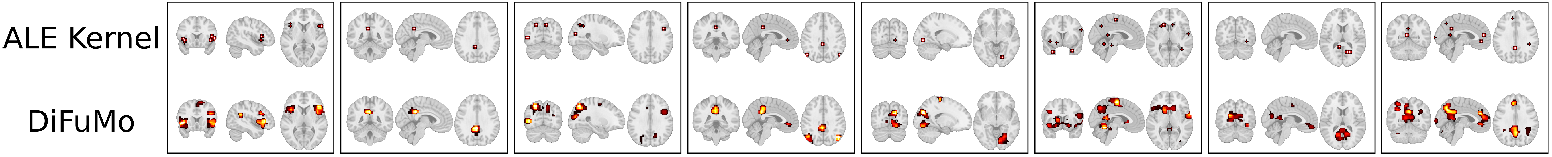
Gaussian products transformed to DiFuMo. Points, from coordinate tables, were first convolved with a Gaussian kernel (top). The image output from Gaussian kernel convolution was reconstructed with DiFuMo (bottom).

### 6.2 Text

Text included titles and abstracts (n=30826) from PubMed publications. All text is paired to a DOI. Text was embedded using Specter2 [26]. This produces a 768-dimensional vector per publication. Specter2 was trained with contrastive loss, promoting ranking similarity between pairs. These embeddings almost collapse to a mean. This results in high cosine similarity for unrelated pairs, e.g. the expected cosine similarity distribution is shifted upwards by a constant. For text-to-brain decoding, the embeddings are demeaned with an empty text embedding, shifting the expected value of cosine similarity between random pairs to zero. Additionally, text embeddings were unit normalized, as contrastive loss is scale insensitive.

### 6.3 NeuroVault

NeuroVault images (*n* = 4553) span several group-level map types reflecting different analysis outputs, including inferential statistics maps: t-, z-, and F-maps (voxelwise test statistics for a contrast model) and p-maps (the corresponding voxelwise significance values). We also included beta maps (univariate GLM estimates and multivariate/decoding weight maps) as well as ROI/masks (binary/probabilistic inclusion maps) and parcellations (discrete region labels), which define spatial subsets or atlas assignments rather than continuous statistics.

Activation maps were cluster-thresholded. For each map, we applied a high-intensity threshold at the 99th percentile and removed small clusters using a minimum cluster extent of 50 voxels. The surviving suprathreshold clusters were then revectorized with the same masker into a fixed-length feature vector, *d* = 28542, and used in testing. Finally, we binarized the resulting vectors (presence/absence of surviving voxels) because the absolute scaling of values across map types (e.g., t-, z-, p-, F-, and beta maps) is not directly comparable; binarization yields a uniform support that emphasizes spatial localization over intensity.

NeuroVault text was limited to titles and abstracts, since per-image contrast definitions were often noisy or missing. Since multiple images were paired to each title-abstract pair, the contrastive model was used to select the most similar image. This inflates scores and provides an upper performance bound. All *n* = 4553 images were used when evaluating the autoencoder, since the autoencoder does not require text-image pairs.

**Table 1.** Autoencoder architecture (layer-wise).

| Layer | Module | Input → Output | Activation |
| --- | --- | --- | --- |
| Encoder 1 | Linear | 28542 → 1024 | ReLU |
| Encoder 2 | Linear | 1024 → 512 | ReLU |
| Encoder 3 | Linear | 512 → 384 | None |
| Decoder 1 | Linear | 384 → 512 | ReLU |
| Decoder 2 | Linear | 512 → 1024 | ReLU |
| Decoder 3 | Linear | 1024 → 28542 | None |

Shapes omit batch dimension.

## 7 Autoencoder

### 7.1 Architecture

After resampling to 4 mm^3^ voxel space, masking, and flattening, each neuroimage is a 28,542 dimensional vector. The exact layers, sizes, and activation functions are provided in Tab. 1. The encoder maps from 28,542 to 384, then the decoder maps from 384 back to 28,542. ReLU activations are used. The model can be thought of as a single masked convolution per layer, with a kernel size equal to the input size. This simplifies handling sparsity from outside-of-brain voxels. Alternatively, convolutional and transposed convolutional layers with a smaller kernel could be explored in future work.

### 7.2 Binary Cross-Entropy Loss

The autoencoder was trained with binary cross-entropy (BCE)-with-logits loss. Let **x**^(*i*)^ ∈ [0, 1]^*D*^ denote a masked and flattened activation map with *d* = 28542. The decoder produces logits **z**^(*i*)^ ∈ ℝ ^*d*^, that are transformed to probabilities using the sigmoid function, *σ*(*u*).

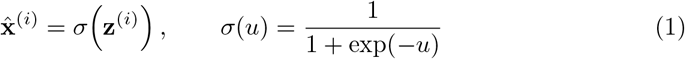

We minimize the binary cross-entropy between **x**^(*i*)^ and 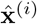 using the numerically stable BCE-with-logits form:

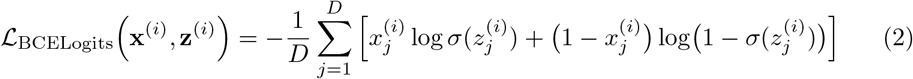

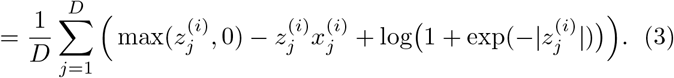

The overall reconstruction loss is the minibatch average over *B* examples:

**Table 2.** Projection head architecture (layer-wise).

| Layer | Module | Input → Output | Activation |
| --- | --- | --- | --- |
| Text 1 | Linear | 768 → 512 | ReLU |
| Text 2 | Linear | 512 → 384 | None |
| Neuro 1 | Linear | 384 → 384 | ReLU |
| Neuro 2 | Linear | 384 → 384 | None |

Shapes omit batch dimension.

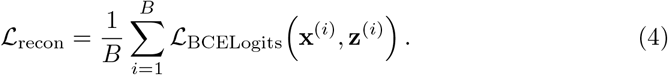

### 7.3 Training

The autoencoder was trained for 100 epochs using AdamW with a learning rate of 5 × 10^−5^ and a batch size of 256. Reconstruction was optimized with binary cross-entropy loss with logits as described above.

## 8 Projection Head

### 8.1 Architecture

There are three types of projection heads. The first two are used to transform 768 dimensional Specter2 embeddings to 384 dimensions. These two projection heads were the same size but trained on different targets. One was trained to map latent text to the autoencoder’s latent space, providing text-to-brain decoding. The other text projection head was learned alongside the neuro projection head, optimized with the contrastive objective, InfoNCE loss. The neuro projection head maps from the autoencoder latent space to the bi-modal latent space. Tab. 2 contains the exact layers sizes.

### 8.2 InfoNCE Loss

Contrastive projection heads were trained with Info Noise Contrastive Estimation (InfoNCE) loss. Let **I** ∈ **ℝ**^*B*×*d*^ and **T** ∈ **ℝ**^*B*×*d*^ represent a batch of image and text embeddings, respectively, where *B* is the batch size and *d* is the embedding dimension. We used *B* = 2048 and *ℓ*_2_-normalize each embedding for *i* ∈ {1, …, *B*}:

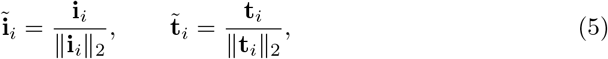

We then compute the pairwise cosine similarity matrix:

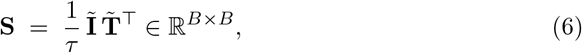

The temperature parameter was learnable and initialized to *τ* = 0.07 [15]. The correct text-image pair for each index *i* is the diagonal entry *S*_*ii*_, all off-diagonal entries are mismatched pairs, i.e., text-image pairs from mismatched studies. Thus, target labels are the identity matrix. We optimize a symmetrized cross-entropy loss over image-to-text (ℒ _i2t_) and text-to-image (ℒ _t2i_) retrieval. The final loss is the mean of the two.

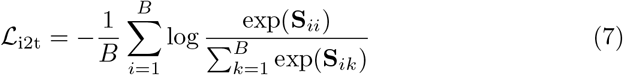

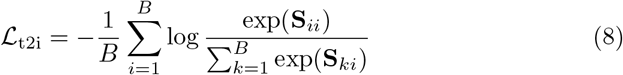

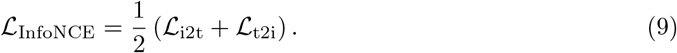

### 8.3 Training

The projection head for text-to-brain generation was trained for 200 epochs with a batch size of 1024 using AdamW with a learning rate of 5 × 10^−5^. Optimization of this text-to-brain model used a BCE-with-logits objective. For joint text–image contrastive alignment, we trained paired projection heads using a symmetric InfoNCE contrastive objective with temperature, *τ* = 0.07. Models were optimized with AdamW for 300 epochs using a batch size of 2048 and learning rate of 1 × 10^−5^.

## 9 Query Transformer

The Qformer maps raw (*r*) and semantic (*s*) latent image embeddings to the token embedding space of the language model, here, the neuroscience-adapted Qwen3-0.6B. The semantic embeddings are raw image embeddings projected onto the text-semantic space using the contrastive projection heads. The language model predicts text, given these latent representations:

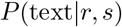

The Qformer transforms the latent embeddings to the token embedding matrix (*Z*) of the LLM. During training, *s* is used. At inference *s*_proj_ may alternatively be used.

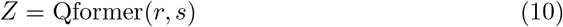

Semantic embeddings may optionally be projected onto a canonical semantic space. This improves robustness to noise and preprocessing variation by replacing a predicted semantic embedding with a soft mixture of clean, interpretable canonical prototypes. Let *s* denote the predicted semantic embedding and let *C* denote a bank of *k* canonical embeddings in the same *d*-dimensional contrastive space:

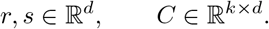

Both *s* and the rows of *C* are *ℓ*_2_-normalized. Projecting *s* onto *C* is the cosine similarity between *s* and each canonical prototype. These similarities are passed through softmax with a temperature parameter (*τ*) that determines smoothing across concept embeddings. With lower temperature, the projection behaves more like nearest-neighbor assignment. With higher temperature, it averages across more canonical concepts. This improves the decoding of maps that are similar enough to, yet outside of, the training distribution.

**Table 3.**
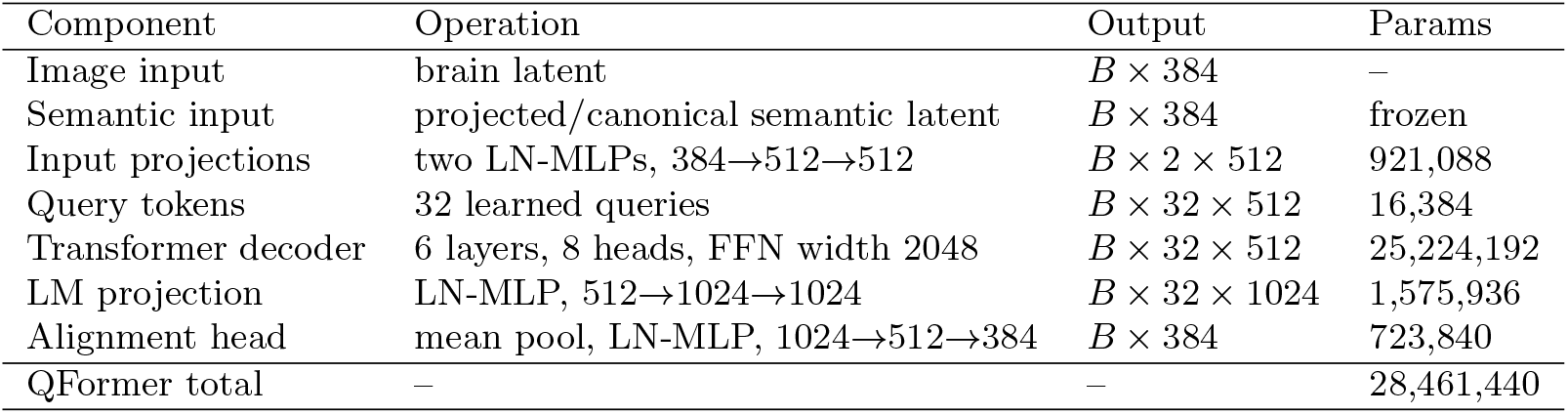
Query transformer architecture. *B* denotes batch size, LN denotes layer normalization, MLP denotes multilayer perceptron, FFN denotes feed-forward network, and LM denotes language model. The packaged model also includes a frozen 295,680-parameter image projection head and frozen canonical semantic projection matrix.

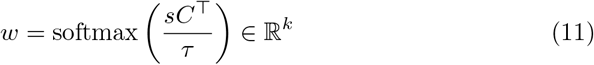

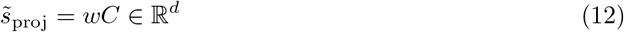

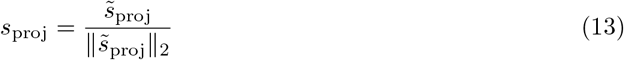

The canonical embedding matrix can be split into subtypes: regions, networks, and function. This allows steering the LLM towards these targets. For example appending the `[NETWORK]` token to the bottom of *Z* steers language generation towards network-level descriptions while `[FUNCTION]` token steers towards higher-level, cognitive interpretations. Appending `[REGION]` generates text related to the most dominant region in the image. Future work could increase these steering tokens to various sub-domains, given curated datasets.

## 10 Generative Performance

**Table 4.** Match-versus-null evaluation of brain-to-text and text-to-brain generation on the PubMed test set (*n* = 2987). Null scores were computed from mismatched image-text pairs. Welch’s *t*-tests compare matched and null distributions; *p*-values are reported in scientific notation.

| Direction | Metric | Match mean | Null mean | $t$ | $p$ |
| --- | --- | --- | --- | --- | --- |
| Brain-to-text | Contrastive cosine | 0.27 | 0.02 | 86.54 | $1.61 \times 10^{-1025}$ |
| Brain-to-text | BERTScore F1 | 0.56 | 0.52 | 68.03 | $8.44 \times 10^{-737}$ |
| Brain-to-text | Semantic cosine | 0.47 | 0.40 | 22.94 | $1.51 \times 10^{-111}$ |
| Text-to-brain | Contrastive cosine | 0.28 | 0.11 | 52.00 | $3.22 \times 10^{-482}$ |
| Text-to-brain | Dice | 0.47 | 0.40 | 24.07 | $3.25 \times 10^{-122}$ |
| Text-to-brain | Pearson | 0.29 | 0.17 | 26.66 | $7.46 \times 10^{-148}$ |

## 11 Model Comparison

**Fig. S3.**
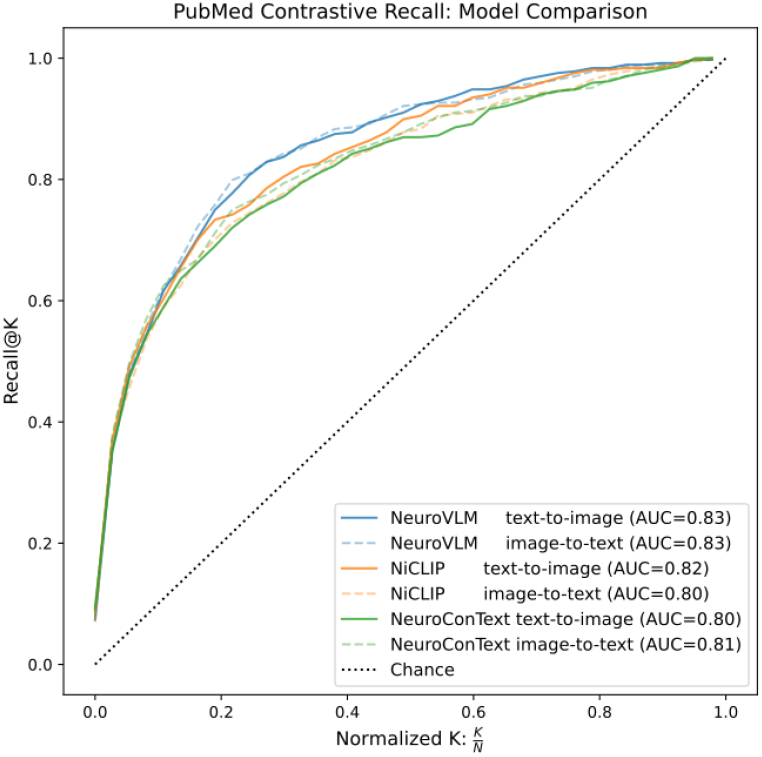
Contrastive model comparison. Recall@k test set curves for NeuroVLM (blue), NiCLIP (orange), and NeuroConText (green). Models performed similarly across the methods. Text includes titles and abstracts only. NeuroVLM uses a language encoder that is 63 times smaller than NiCLIP and NeuroConText, 110 million versus 7 billion parameters.

## 12 Independent Component Labeling

**Fig. S4.**
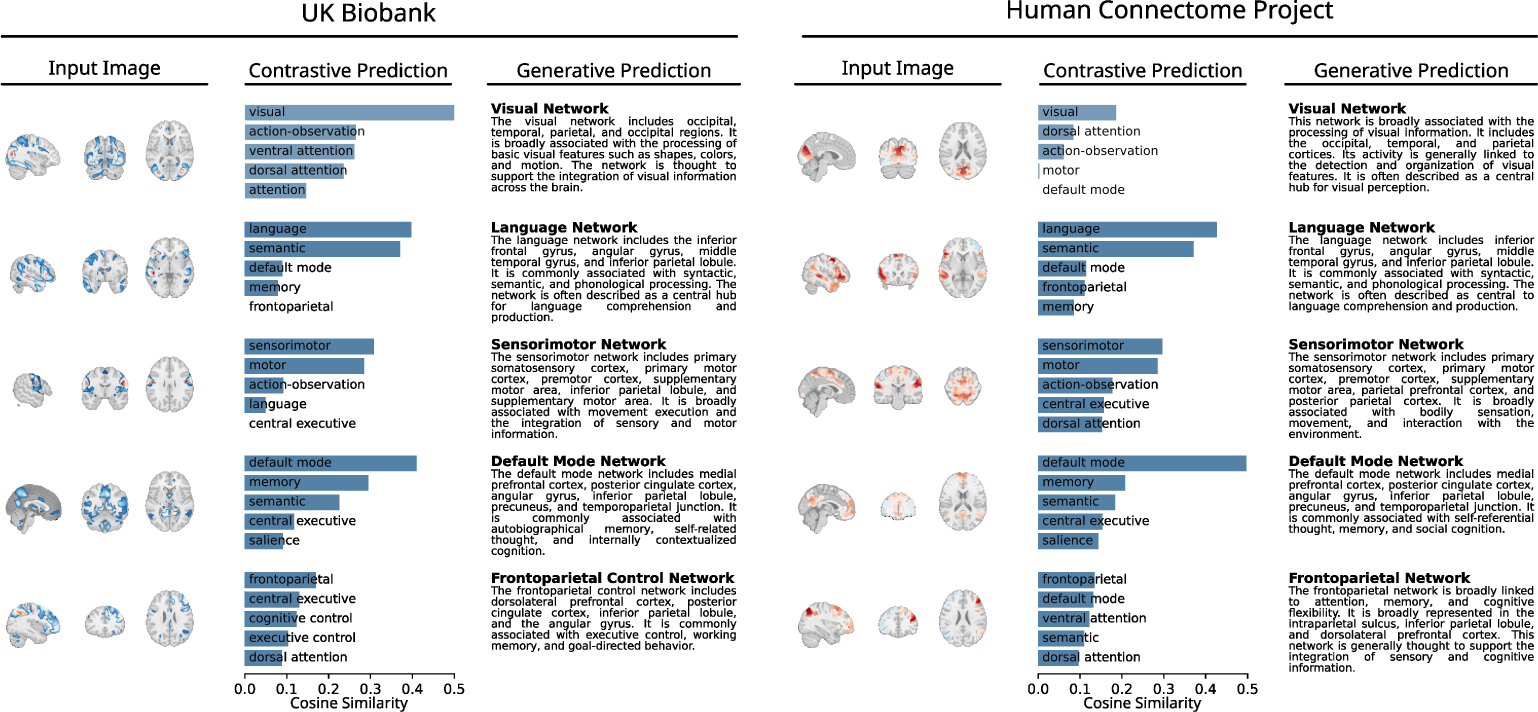
Independent Component Labeling. Automated labeling of resting-state components from the Human Connectome Project and UK Biobank. Each row corresponds to one network. The contrastive columns are the top-5 most similar pairs from the contrastive models, taking argmax gives the predicted label for that image. The generated column is brain-to-text generation using the Qformer model.

